# Oculomotor capture reveals trial-by-trial neural correlates of attentional guidance by contents of visual working memory

**DOI:** 10.1101/320259

**Authors:** Valerie M. Beck, Timothy J. Vickery

## Abstract

Evidence from attentional and oculomotor capture, contingent capture, and other paradigms suggests that mechanisms supporting human visual working memory (VWM) and visual attention are intertwined. Features held in VWM bias guidance toward matching items even when those features are task irrelevant. However, the neural basis of this interaction is underspecified. Prior examinations using fMRI have primarily relied on coarse comparisons across experimental conditions that produce varying amounts of capture. To examine the neural dynamics of attentional capture on a trial-by-trial basis, we applied an oculomotor paradigm that produced discrete measures of capture. On each trial, subjects were shown a memory item, followed by a blank retention interval, then a saccade target that appeared to the left or right. On some trials, an irrelevant distractor appeared above or below fixation. Once the saccade target was fixated, subjects completed a forced-choice memory test. Critically, either the target or distractor could match the feature held in VWM. Although task irrelevant, this manipulation produced differences in behavior: participants were more likely to saccade first to an irrelevant VWM-matching distractor compared with a non-matching distractor – providing a discrete measure of capture. We replicated this finding while recording eye movements and scanning participants’ brains using fMRI. To examine the neural basis of oculomotor capture, we separately modeled the retention interval for capture and non-capture trials within the distractor-match condition. We found that frontal activity, including anterior cingulate cortex and superior frontal gyrus regions, differentially predicted subsequent oculomotor capture by a memory-matching distractor. Other regions previously implicated as involved in attentional capture by VWM-matching items showed no differential activity across capture and no-capture trials, even at a liberal threshold. Our findings demonstrate the power of trial-by-trial analyses of oculomotor capture as a means to examine the underlying relationship between VWM and attentional guidance systems.

## 1 Introduction

Visual attention and visual working memory (VWM) are deeply intertwined and interactive cognitive systems (Chun, 2011; Desimone, 1996; Gazzaley & Nobre, 2012; Kiyonaga & Egner, 2013). Some claim that attention is automatically captured by VWM-matching objects (Folk, Remington, & Johnston, 1992; Soto, Hodsoll, Rotshtein, & Humphreys, 2008), while others disagree (Downing & Dodds, 2004; Houtkamp & Roelfsema, 2006). This ongoing controversy raises a fundamental question regarding the interface between attention and VWM – when and why do VWM representations obligatorily influence attention? Understanding the neural substrate of this interaction between visual attention and VWM is imperative for resolving such persistent debates in the literature and advancing a unified theory of attention and working memory.

Prior work examining the neural correlates of attentional capture has identified frontal and parietal brain regions whose activity is correlated with behavioral interference from a singleton distractor (de Fockert, Rees, Frith, & Lavie, 2004) or enhanced memory for salient stimuli (Wills et al., 2016). Using a variation of the additional singleton paradigm (Theeuwes, 1992), de Fockert et al. (2004) found increased activity in frontal (left lateral precentral gyrus) and parietal (bilateral superior parietal lobules) regions when a salient distractor was present versus absent from the search array. Furthermore, increased frontal cortex activity predicted decreased interference from the salient singleton distractor suggesting that increased cognitive control mitigated the behavioral impact of the salient distractor. Extending this work to contingent attentional capture (Folk et al., 1992), Wills et al. (2016) used an RSVP (rapid serial visual presentation) task to identify brain regions whose activity was correlated with attentional capture by a letter displayed in a task-relevant color. Participants were asked to remember a series of letters in the order presented while also monitoring for a pound sign displayed in a prespecified color (e.g., red). Memory accuracy for letter identity when the letter appeared in the task-relevant color (e.g., red) was greater than for letters displayed in a control color. Again, frontal (right inferior frontal junction) and parietal (right superior parietal lobule) regions showed increased activity during encoding of color-matching letters compared to encoding of control letters. Moreover, increased activity in frontal cortex during encoding was correlated with reduced memory accuracy, though this may have been driven by individuals with particularly poor memory performance. In sum, prior work suggests that both frontal and parietal regions, including regions previously associated with dorsal and ventral attention networks (Corbetta & Shulman, 2002) and cognitive control (Harding, Harrison, Breakspear, Pantelis, & Yücel, 2016), may be involved in attentional capture by salient and task-relevant items.

One complicating factor for identifying the neural correlates of attentional guidance toward VWM-matching objects is that attentional “capture” has typically been quantified using continuous measures like manual response times (RTs) that are uninformative at a single trial level. For example, Soto and colleagues (2005) instructed subjects to remember a colored outlined shape, then search for a shape that contained a tilted line among other shapes that contained vertical lines. Subjects reported the direction of the tilted line, then indicated whether a memory probe was the same or different from the item presented at the beginning of the trial. Subjects had slower RTs when the VWM-matching distractor contained a vertical line (invalid condition) than when it was not present in the array (neutral condition) and had faster RTs when the VWM-matching item had a tilted line and was the target item (valid condition). Slowed RTs in the invalid relative to the neutral condition suggest that attention was automatically captured by the VWM-matching distractor, regardless of whether it was relevant to the task. However, because a single RT does not indicate whether attentional capture occurred on any particular trial, the invalid condition necessarily contained trials in which attention was captured by a VWM-matching item, and also very likely included trials when the VWM-matching item did not capture attention.

The possibility that VWM-matching distractor trials sometimes do and sometimes do not result in capture raises questions about later studies that used this same paradigm during fMRI scanning to identify the neural mechanisms of VWM-based attentional guidance (Soto, Greene, Chaudhary, & Rotshtein, 2012; Soto, Greene, Kiyonaga, Rosenthal, & Egner, 2012; Soto, Humphreys, & Rotshtein, 2007). Different patterns of neural activity were identified using various contrasts (valid > invalid; valid + invalid > neutral; valid + invalid > repetition detection), but none of these strictly isolated instances of VWM-based attentional guidance to compare against instances when VWM contents do not influence attentional guidance. In addition to the fact that these conditions may not perfectly correspond to VWM-based guidance, the composition of the displays used in the valid, invalid, and neutral conditions necessarily differ in terms of whether the target item (tilted line) and the memory-matching shape occur at the same location (valid trials), different locations (invalid trials), or are not both present (neutral trials), which further complicates interpretation of the results. Finally, because the index of attentional capture in these studies was based on RT differences, the contrasts are necessarily confounded with RT. Many regions of the brain are known to be generally covary with RT differences (Yarkoni, Barch, Gray, Conturo, & Braver, 2009), and some prior findings may in part reflect such covariation rather than indicating the correlates of VWM-based attentional capture. More broadly, these contrasts may have indexed neural activity both underlying and caused by capture.

Studies applying this paradigm have yielded inconsistent results. Soto et al. (2007) identified a network of brain regions (including the parahippocampal gyrus, lingual gyrus, and superior frontal gyrus) that respond to a VWM-matching item distinct from the reappearance of an item not held in VWM. Later studies (Soto et al., 2011; Soto, Greene, Chaudhary, et al., 2012) yielded greater activity in bilateral occipital regions with little (Soto et al., 2011) or no (Soto, Greene, Chaudhary, et al., 2012) response from the superior frontal gyrus region. The complications listed above may have contributed to inconsistencies in resulting outcomes. Thus, we sought converging evidence using a different paradigm that clearly dissociates capture and non-capture trials, while holding equal as many other factors as possible.

In a combined memory and eye movement task (see Figure 1), participants are frequently “captured” by a memory-matching distractor (T_N_D_M_ condition) and make an eye movement to that item before proceeding on to the saccade target. Recent work using this “Remote Distractor” paradigm to compare oculomotor capture by a VWM-matching distractor under high and low VWM loads demonstrates the power of using oculomotor measures as opposed to traditional RT measures (Beck & Vickery, *submitted*). Oculomotor behavior revealed significantly greater capture by a VWM-matching distractor compared to a non-matching distractor under both a memory load of one item and two items. Importantly, the observed capture was significantly reduced for load-2 compared to load-1 trials, suggesting that multiple VWM representations were held in a mixture of active (able to influence attentional guidance) and accessory (unable to influence attentional guidance) states. However, a time-to-target-fixation (analogous to RT) analysis on the same data revealed significantly greater capture by a VWM-matching distractor on load-1, but *not* load-2 trials. This discrepancy between oculomotor and time-to-target-fixation results illustrates the value of discrete measures of capture such as the one used here. By recording eye movements while participants performed this type of task in the scanner, we can use this oculomotor measure (saccade to the distractor) to directly compare *capture* trials – when VWM interacted with attentional guidance – against *non-capture* trials – when VWM did *not* interact with attentional guidance. Comparing neural activity based on discrete behavioral outcomes while under the same task conditions and the same visual displays (unlike previous studies) allows better isolation of the neural mechanisms that support interaction between visual attention and VWM. Previewing our findings, we discovered a much more constrained set of regions, including superior frontal gyrus (SFG) and anterior cingulate cortex (ACC), whose activity during the memory retention interval was positively correlated with attentional capture by VWM-matching distractors.

**Figure 1:**
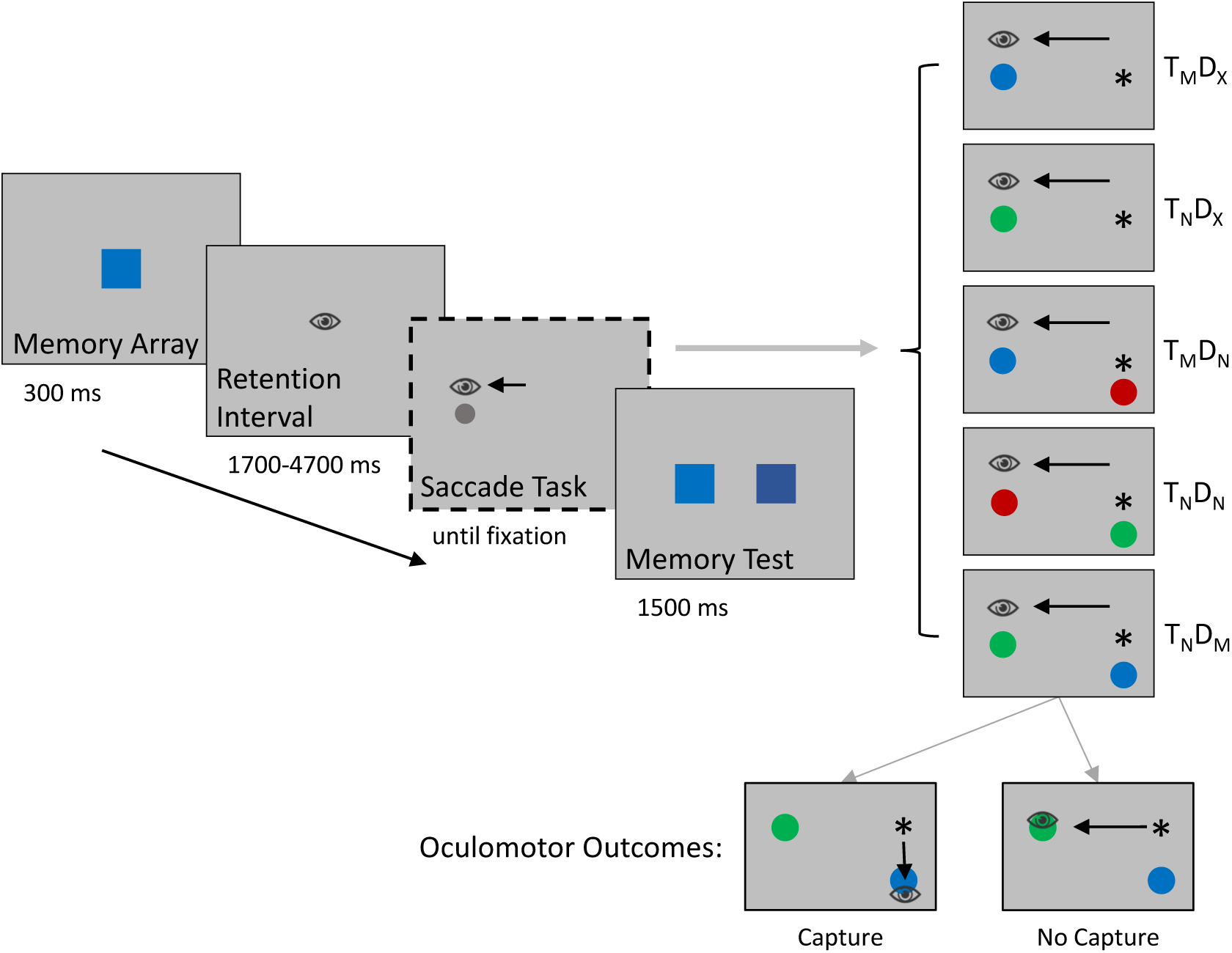
Illustration of trial events, trial types (T = target, D = distractor, M = match, N = non-match, X = absent), and discrete oculomotor outcomes (“capture”, “no capture”). Participants were instructed to remember the memory array color, make an eye movement to a left or right target disk, then indicate which of the two colors presented during the memory test matched a color in memory. The saccade task yielded five different trial types, depending on whether the target or distractor disks matched a color in memory. All trial types were equally likely and randomly intermixed within each block.

**Figure 2:**
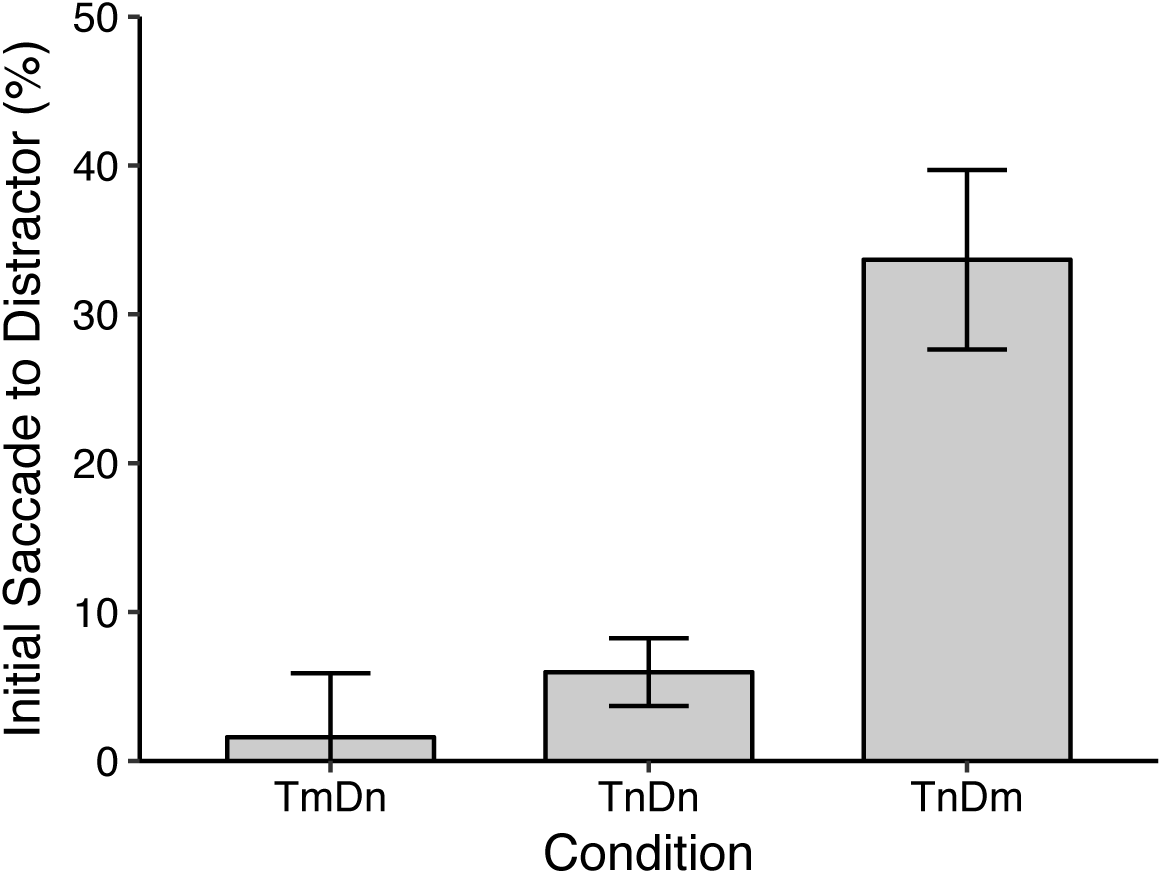
Initial saccades were more frequently directed to a memory-matching distractor compared to a non-matching distractor [34% vs. 6%; t(14) = 8.27, p < .001]. Error bars indicate +/- within-subject 95% confidence intervals (Morey, 2008).

## 2 Method

### 2.1 Participants

Eighteen University of Delaware students who were 18-40 years of age (15 Female) and reported having normal color vision were compensated $20/hour for one 2-hour session. All participants were right-handed and reported no known psychological or neurological conditions and no current use of psychoactive medications. All procedures were approved by the University of Delaware Institutional Review Board.

### 2.2 Stimuli, Apparatus, and Procedure

Stimuli were presented on a 32” BOLDscreen LCD monitor (120 Hz) placed at the end of the bore of the scanner (approximate viewing distance of 150 cm). Eye position was recorded at 1000 Hz using an Eyelink 1000 long-range eye-tracking system. Saccades were defined using a combined velocity (>35°/s) and acceleration (>9500°/s^2^) threshold. To ensure accurate eye movement recording, participants completed a 9-point calibration at the beginning of each block (every 40 trials).

As illustrated in Figure 1, each trial began with a 300-ms presentation of a colored square, subtending 1.6 degrees visual angle (hereafter dva), positioned at central fixation. After a blank retention interval (1700, 2700, 3700, or 4700 ms; uniform distribution, 3200 ms average duration), a target disk (0.86 dva) appeared to the left or right of fixation (the center of the disk was positioned at 4.61-7.06 dva from fixation). The target disk was equally likely to match or not match the memory color. On some trials, the left/right target disk appeared simultaneously with a distractor disk (0.94 dva) located above or below fixation (2.14 dva from fixation). The distractor disk could also match the memory color yielding five trial types (see Figure 1): target match, no distractor (T_M_D_X_); target non-match, no distractor (T_N_D_X_); target match, distractor non-match (T_M_D_N_); target non-match, distractor non-match (T_N_D_N_); and target non-match, distractor match (T_N_D_M_). Participants were instructed to make an eye movement to the left/right target disk and ignore up/down distractor disks. The target disk and distractor disk (if present) remained on the screen for 500 ms. Once the saccade task was completed, there was a 500-ms blank delay, then two squares (1.2 dva each) appeared on either side of fixation (offset by 2.0 dva) and participants were instructed to indicate via key press which of the two squares matched the color in memory (left button for left square match, right button for right square match). One square matched the color presented at the beginning of the trial, while the other was a foil drawn from the same color family. The memory test remained on screen for 1500 ms, then feedback (correct/incorrect) was displayed for 500 ms.

Object colors were drawn from four different color families (reds, greens, blues, or pinks) that each contained four exemplars, and objects were presented on a light gray background. The memory array color was selected randomly for each trial (e.g., light blue) and the memory test foil was another color from the same family (e.g., navy blue). Colors of any non-matching target or distractor disks were selected from the remaining color families (e.g., reds, greens).

Participants completed 40 practice trials once they were situated in the scanner and then completed 10 blocks of experimental trials with a short break in between blocks. Each block contained 40 total trials, 8 each of the 5 trial types. Trials from each condition were randomly intermixed within each block.

### 2.3 Imaging Procedure

Structural and functional data were acquired on a 3T Siemens Prisma system with a 64-channel head/neck coil. We acquired a high-resolution (0.7 mm isometric voxels) T1-weighted MPRAGE structural image that was used for anatomical reconstruction and participant coregistration. Functional scans were T2*-weighted images collected using a multi-band echoplanar imaging sequence consisting of 66 slices with an oblique axial orientation (approximately 25° from AC-PC alignment) and acquired with a resolution of 2.2 mm × 2.2 mm × 2.2. mm (sequence parameters: TR=1 s, TE = 39.4 ms, flip angle 90°). Ten functional runs consisting of 334 volumes (plus 4 initial acquisitions which were automatically discarded) were acquired for each participant, with each run lasting 5 minutes, 34 seconds.

### 2.4 Structural and Functional Preprocessing

MRI analyses were conducted using fMRIB Software Library (FSL; FMRIB’s Software Library, www.fmrib.ox.ac.uk/fsl) version 5.0.9 and FMRI Expert Analysis Tool (FEAT) version 6.0 (Jenkinson, Beckmann, Behrens, Woolrich, & Smith, 2012). Structural scans were skull-stripped using BET (Smith, 2002) and registered to a standard MNI152 2-mm template. Prior to statistical analysis, each functional run was subjected to motion correction, high-pass temporal filtering (100 s cutoff), skull stripping, alignment to high-resolution structural images, and smoothing with an 8-mm full width at half maximum (FWHM) Gaussian kernel. fMRI data were analyzed in three hierarchical levels.

#### 2.4.1 First-level analysis

At the first level, each run was modeled using a standard GLM approach with 7 base regressors, each of which was convolved with a double-gamma standard hemodynamic response function prior to entry into the model. Each of the 7 base regressors were accompanied by their first temporal derivatives to improve fit. Three unit-height boxcar regressors modeled the memory array (1 s duration), saccade task (1 s duration), and memory test and feedback (2 s duration) periods. The most critical regressors were those that modeled the retention interval – time between appearance of the memory array and appearance of the saccade target. Although the total duration of this retention interval varied (1.7-4.7 s), the regressors modeled the final 1 second preceding appearance of the saccade target.

For the distractor-match (T_N_D_M_) condition, trials were categorized based on whether the initial eye movement (see 2.5 “Eye Movement Analysis” for details) went to the memory-matching distractor (“capture”) or went straight to the saccade target (“non-capture”). Because the eye movement data collected in the scanner was noisier than data from a typical desktop eye-tracking system and in order to maximize power for the fMRI analyses, eye movements from distractor-match trials (T_N_D_M_) were inspected individually by one of the authors (VMB) and classified as “capture” or “no capture” based on the latency and trajectory of the first 1-2 eye movements after the saccade target and distractor disks appeared. This resulted in 5.4% of trials that were reclassified from “no capture” to “capture” or vice versa. For each run, the number of “capture” and “non-capture” trials included in the first-level analysis were equated, and excess trials were added to a junk regressor.

The retention interval for all remaining conditions (T_M_D_X_, T_N_D_X_, T_M_D_N_, T_N_D_N_) was modeled in a separate regressor. A contrast was defined at the first level to determine which voxels whose activity significantly predicted subsequent oculomotor capture or non-capture by a memory-matching distractor.

#### 2.4.2 Second-level analysis

Contrast of parameter estimate images (COPEs) from the first level were converted to standard space (MNI152, 2-mm), and then combined across runs completed by each subject in a second-level, fixed-effects analysis. Runs that did not contain at least one “capture” trial for the distractor-match (T_N_D_M_) condition were excluded from analysis.

#### 2.4.3 Third-level analysis

COPEs from the second-level were combined across subjects in a third-level, mixed-effects analysis. Clusters were defined using a family-wise error (FWE) correction following a Z > 2.8 threshold (*p* < .005), based on Gaussian Random Field theory.

### 2.5 Eye Movement Analysis

Analysis of eye movement data focused on the trajectory and latency of the first eye movement that occurred after the saccade target and distractor (if present) disks appeared. These saccades were categorized as being directed either toward the target (left/right horizontal meridian ±45°) or toward the distractor (upper/lower vertical meridian ±45°). Only trials during which the first eye movement was directed to a relevant location (up/down/left/right 90° wedge containing a target or distractor disk) were included for analysis [8.4% of all trials rejected]. Trials with eye movements that were excessively fast (< 90 ms) or slow (> 600 ms) [9.8% of all trials rejected] or that were missing eye movement data [7.2% of all trials rejected] were excluded. For all remaining trials [88.5% of trials retained (some trials met multiple exclusion criteria)], when an initial eye movement was directed toward a distractor, the trial was categorized as a “capture” trial whereas if the initial eye movement was directed toward the target, the trial was categorized as “no capture” (see Figure 1 for example).

## 3 Results

Participants who executed eye movements to a distractor location on 10% or more of distractor-absent trials, indicative of poor eye tracking or misunderstanding of the task, were excluded from analysis (N=1). Additionally, participants who had fewer than 10 total “capture” trials were excluded from analysis (N=2). The following analyses are based on data from the remaining participants (N=15, 12 Female).

### 3.1.1 Memory accuracy

To test for differences in memory accuracy across trial types, memory test manual response accuracy was entered into a one-way 5-level (trial type: T_M_D_X_, T_N_D_X_, T_M_D_N_, T_N_D_N_, T_N_D_M_) repeated-measures ANOVA. There was a significant effect of trial type [*F*(4, 56) = 4.89, *p* = .002, 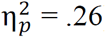] with slightly greater memory test accuracy when the saccade target matched [T_M_D_X_: M = 93.9%; T_M_D_N_: M = 92.5%] than when it did not match [T_N_D_X_: M = 88.6%, *t*(14) = 2.67, *p* = .018; T_N_D_N_: M = 87.9%, *t*(14) = 2.73, *p* = .016]. Memory test accuracy was marginally greater when the distractor matched the item in memory than when it did not [T_N_D_M_: M = 91.8%, *t*(14) = 2.06, *p* = 0.06], but there was no difference in memory test performance when participants made an eye movement to the memory-matching distractor (M = 92.4%) or did not (M = 92.0%). Overall, memory test performance was uniformly high across conditions, suggesting that the memory-matching distractor could not aid performance if attended.

### 3.1.2 Oculomotor capture

The critical analysis for the current study examined the probability that an eye movement was directed to a memory-matching distractor. For this, we examined the trajectory of the first eye movement that occurred after the saccade target and distractor (if present) disks appeared. The proportion of “capture” trials was calculated for each distractor-present condition (see 2.5 “Eye Movement Analysis”). As expected, and consistent with previous studies (Beck & Vickery, submitted; Hollingworth, Matsukura, & Luck, 2013), the probability of being captured by and making an eye movement toward an irrelevant distractor was greater when the distractor matched the memory item color [T_N_D_M_: M = 33.7%] than when it did not [T_N_D_N_: M = 6.0%; *t*(14) = 8.27, *p* < .001].

### 3.1.3 fMRI data

To examine which brain regions were more active during VWM-based attentional guidance, we contrasted retention interval activity preceding instances of “capture” – when VWM contents influence attentional guidance – with instances of “no capture” – when VWM contents do not influence attentional guidance (as described in 2.4.1 “First-level analysis”). Of the regions previously associated with VWM-based attentional guidance (Soto et al., 2007), only activity in superior frontal gyrus (SFG) predicted oculomotor capture by a VWM-matching distractor (see Figure 3 and Table 1). In addition to activity in SFG, we also found elevated activity in anterior cingulate cortex (ACC) that predicted oculomotor capture by a VWM-matching distractor.

**Figure 3:**
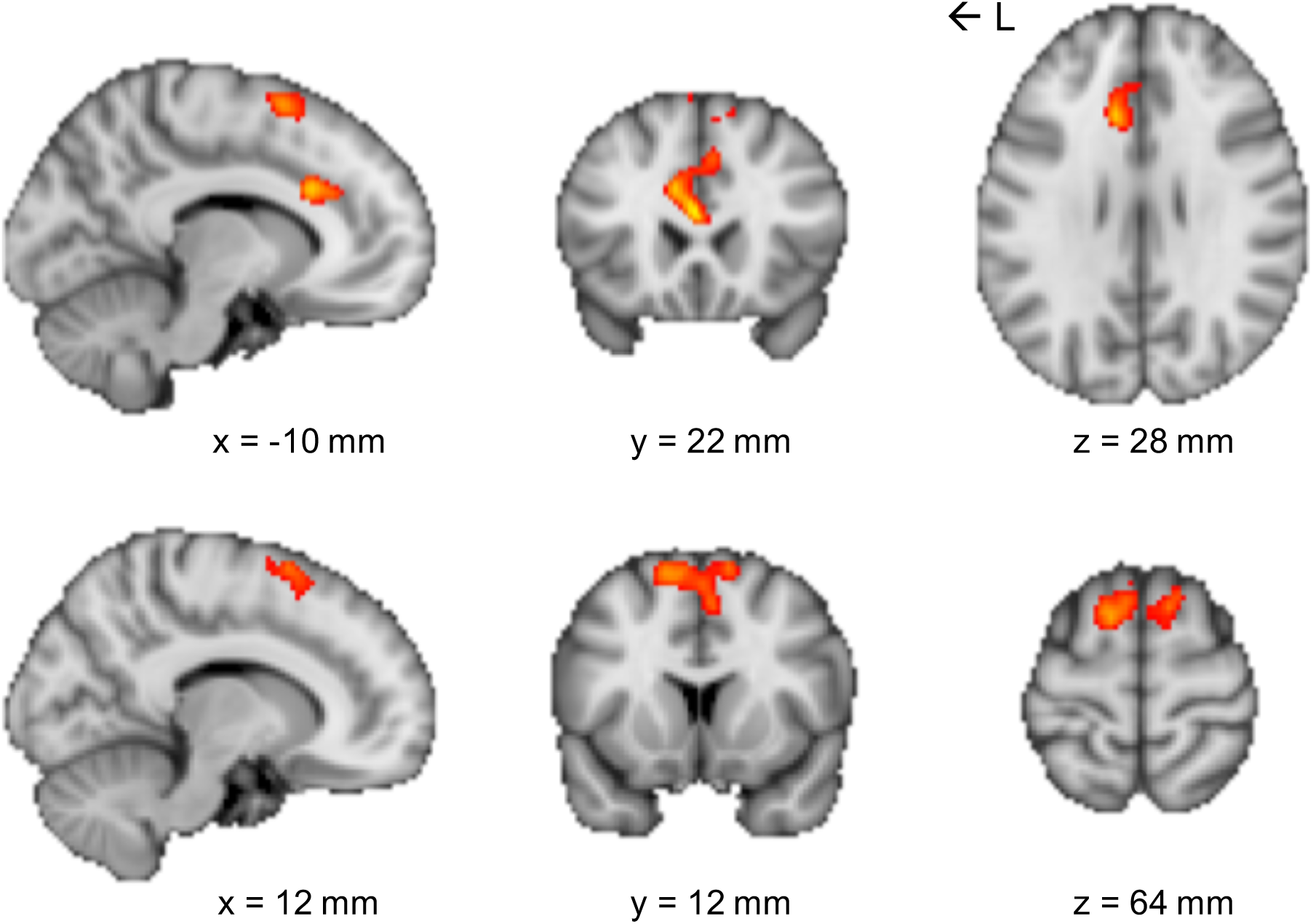
Regions showing an increased response during the last 1 second of the retention interval preceding oculomotor capture by a memory-matching distractor compared to no oculomotor capture. Clusters were defined using FWE correction following a Z > 2.8 threshold. The capture > no capture contrast yielded one cluster that included anterior cingulate cortex (ACC) and superior frontal gyrus (SFG) regions (see Table 1). Top row coordinates are centered on left ACC, bottom row coordinates are centered on right SFG. Coordinates are in MNI standard space.

**Table 1:**
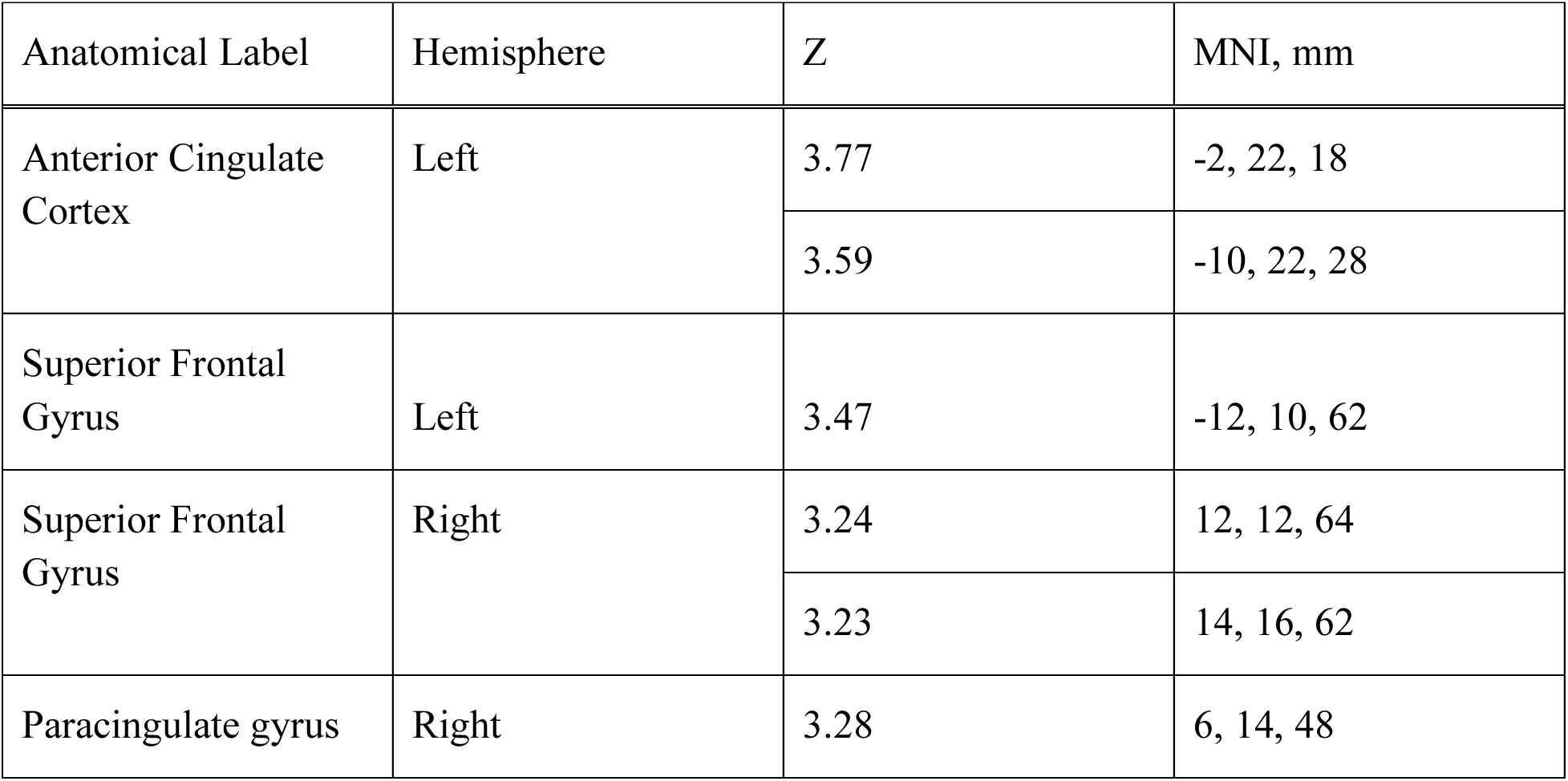
Local maxima for the single large cluster of activation during the retention interval that predicted oculomotor capture by a VWM-matching distractor.

To more explicitly test whether regions previously identified may also predict oculomotor capture by a VWM-matching distractor, we created 8-mm spherical regions-of-interest (ROIs) centered on peak coordinates from previously reported brain regions that showed VWM-modulated activation (Soto et al., 2007). We created ROIs for left and right SFG, left supramarginal gyrus, left and right parahippocampal gyrus, and left lingual gyrus (see Table 2). BOLD activity from each ROI was averaged across participants and tested whether it was significantly different from zero. None of the ROIs yielded activity that reliably predicted oculomotor capture by a VWM-matching distractor (all *p*s > .11) suggesting that the neural correlates of VWM-based attentional guidance identified using the current oculomotor paradigm are distinct from those previously identified using a different paradigm that relied on manual response times.

**Table 2:** Coordinates for brain regions that previously demonstrated VWM-modulated activation (Soto et al., 2007). These coordinates were used to create spherical ROIs for the current analysis.

| Anatomical Label | Hemisphere | MNI, mm | Capture vs. No-Capture contrast |
| --- | --- | --- | --- |
| Superior frontal gyrus (SFG) | Left | -24, 18, 45 | $t(14) = 1.72, p = .108$ |
| | Right | 18, 30, 48 | $t(14) = 0.20, p = .841$ |
| Supramarginal gyrus (SMG) | Left | -42, -36, 60 | $t(14) = -0.16, p = .877$ |
| Parahippocampal gyrus (PHG) | Left | -15, -36, -3 | $t(14) = 0.59, p = .564$ |
| | Right | 24, -42, -6 | $t(14) = 0.84, p = .416$ |
| Lingual gyrus (Ling G) | Left | -3, -72, -9 | $t(14) = -0.45, p = .657$ |

## 4 Discussion

Using a paradigm that better isolates instances of interaction between VWM and visual attention, we found that increased activity in ACC and SFG brain regions during the memory retention interval reliably predicted oculomotor capture by a memory-matching distractor. Both ACC and SFG have previously been implicated in several cognitive processes, particularly higher-level aspects of cognition like executive control, error monitoring, and top-down control. Using a target template or feature (e.g., blue) to guide allocation of attention is a classic example of “top-down” or goal-driven attentional control, so it unsurprising that we would observe increased activity in brain regions previously associated with these processes, even though the memory color in the current paradigm was task-irrelevant.

This pattern of neural activation differs from that previously found using a paradigm that relied on RT measures of VWM-based attentional capture (Soto et al., 2007). Unlike the contrasts formed in those studies, the conditions underlying our primary contrast were equated in terms of the stimulus composition. Furthermore, even though prior work included RT as a covariate (Soto et al., 2007), that may not have completely controlled for systematic differences in RT across conditions and the resultant pattern of activation may at least in part reflect RT-modulated instead of VWM-modulated differences in attentional control. Finally, RT measures necessarily collapse across instances of VWM-based attentional capture and no capture, potentially obscuring the neural correlates of VWM-based attentional guidance. The current paradigm used discrete oculomotor outcomes to identify instances of capture by a VWM-matching distractor, indicating interaction between VWM and attention systems, and instances of non-capture, suggesting lack of interaction between VWM and attention systems. We then compared neural activity preceding instances of capture and non-capture to determine whether different neural states could differentially predict oculomotor outcomes and found that increased activity in ACC and SFG regions reliably predicted instances of capture compared to non-capture.

Networks associated with VWM and attentional control are prevalent throughout the brain, and the specific structures involved tend to vary based on the paradigm used or task demands (Corbetta & Shulman, 2002). More specifically, it has been challenging to identify the structures responsible for supporting the interaction between these cognitive systems. Items held in VWM can be reliably decoded from activity in primary visual cortex (Harrison & Tong, 2009; Serences, Ester, Vogel, & Awh, 2009). These patterns of activation could interact with incoming sensory processing to bias attention toward matching items, a process known as the sensory recruitment hypothesis. If this mechanism had been driving the observed oculomotor capture by VWM-matching distractors, we might expect that regions of occipital cortex would reliably predict the capture and no-capture oculomotor outcomes. Alternatively, and consistent with the current results, the biasing signal driving the interaction between VWM and attention systems could have come from frontal structures previously associated with goal-directed attentional control, such as ACC and SFG. That said, our analysis looked only for elevated activity under capture vs. no-capture, which may obscure visual regional pattern variation that could predict attentional capture. Future work should examine patterns of activation in visual regions during the memory retention interval, and ask whether the strength of sensory recruitment matching the memory item might predict the occurrence of capture.

It has been proposed that representations in VWM can exist in different states – “active” and “accessory” – such that an item in an active state can influence attentional guidance whereas an item in an accessory state cannot (Olivers, Peters, Houtkamp, & Roelfsema, 2011). The observed difference in neural activity preceding capture and no-capture oculomotor outcomes in the current study provides support for this division of VWM representations into active and accessory states. Whether an item in VWM is elevated to an active state or relegated to an accessory state should be determined once the item has been encoded into VWM but before attention can be deployed. It was previously proposed that there may be a frontal gating mechanism that separates VWM representations into active and accessory states (Olivers et al., 2011) and the current results suggest that ACC and/or SFG could be the neural substrate for this gating mechanism.

Fully characterizing the neural substrate for this interaction between VWM and attention systems is important not only for understanding normal visual function but also for understanding and potentially mitigating disordered function. In fact, when patients with schizophrenia (PSZ) were asked to perform the same combined memory and oculomotor task while eye movements were recorded, PSZ more frequently made an initial saccade toward a memory-matching distractor than healthy control subjects (Luck et al., 2014), suggesting an abnormal interaction between VWM and attention systems. Evidence from lesion studies indicates both cortical (Van der Stigchel, van Koningsbruggen, Nijboer, List, & Rafal, 2012) and subcortical (Van der Stigchel, Arend, van Koningsbruggen, & Rafal, 2010) mechanisms for resolving competition between task-relevant and salient distractor objects, but further work is needed to determine whether eye movements toward memory-matching items would be subject to control by the same circuitry.

## 5 Conclusions

In sum, the current work examined the neural signatures of the interaction between attention and VWM by implementing an innovative paradigm that generates a discrete measure of attentional capture. Previous work examining this question has largely relied on continuous measures like manual RTs that are uninformative on a single trial level. Comparing neural activity based on discrete behavioral outcomes while under the same task conditions revealed that activity in ACC and SFG reliably predicted oculomotor capture by a memory-matching distractor. Further work will be needed to determine whether either or both of these regions is necessary or sufficient for supporting the interaction between VWM and attention systems. Elucidating the mechanisms that support the interaction will facilitate progress toward resolving the ongoing debate regarding VWM-based attentional guidance, and further basic understanding and model-building surrounding both attention and VWM.

## Acknowledgements

This research was supported by National Science Foundation grants (BCS 1558535 and OIA 1632849) to Timothy J. Vickery.

